# Vanilloids Hamper *Caenorhabditis elegans* Response to Noxious Heat

**DOI:** 10.1101/2020.09.10.291237

**Authors:** Bruno Nkambeu, Jennifer Ben Salem, Francis Beaudry

**Affiliations:** Groupe de Recherche en Pharmacologie Animal du Québec (GREPAQ), Département de Biomédecine Vétérinaire, Faculté de Médecine Vétérinaire, Université de Montréal, Saint-Hyacinthe, Québec, Canada; Centre de recherche sur le cerveau et l’apprentissage (CIRCA), Université de Montréal, Montréal, Québec, Canada; Institut des Maladies Métaboliques et Cardiovasculaires, INSERM UMR1048, Université de Toulouse, France

**Keywords:** *Caenorhabditis elegans*, Eugenol, Vanillin, Zingerone, Vanilloids, Nociception, Transient receptor potential cation channel

## Abstract

Eugenol, a known vanilloid, was frequently used in dentistry as a local analgesic in addition, antibacterial and neuroprotective effects were also reported. Eugenol, capsaicin and many vanilloids are interacting with the transient receptor potential vanilloid 1 (TRPV1) in mammals and are activated by noxious heat. The pharmacological manipulation of the TRPV1 has been shown to have therapeutic value. *Caenorhabditis elegans* (*C. elegans*) express TRPV orthologs (e.g. OCR-2, OSM-9) and it is a commonly used animal model system to study nociception as it displays a well-defined and reproducible nocifensive behavior. After exposure to vanilloid solutions, *C. elegans* wild type (N2) and mutants were placed on petri dishes divided in quadrants for heat stimulation. Thermal avoidance index was used to phenotype each tested *C. elegans* experimental groups. The results showed that eugenol, vanillin and zingerone can hamper nocifensive response of *C. elegans* to noxious heat (32°C – 35°C) following a sustained exposition. Also, the effect was reversed 6h post exposition. Furthermore, eugenol and vanillin did not target specifically the OCR-2 or OSM-9 but zingerone did specifically target the OCR-2 similarly to capsaicin. Further structural and physicochemical analyses were performed. Key parameters for quantitative structure-property relationships (QSPR), quantitative structure-activity relationships (QSAR) and frontier orbital analyses suggest similarities and dissimilarities amongst the tested vanilloids and capsaicin in accordance with the relative anti-nociceptive effects observed.

## Introduction

*Caenorhabditis elegans* (*C. elegans*) is a commonly used animal model system to study neuronal communication and more recently neurodegenerative diseases [1]. Adult self-fertilizing hermaphrodite *C. elegans* consists of 959 cells, of which 302 are neurons, making this model very attractive to study nociception at the physiological level [2]. *C. elegans* males develop rarely (~ 0.1%). Likewise, *C. elegans* is especially useful to study nociception as it displays a well-defined and reproducible nocifensive behavior, involving a reversal and change in direction away from the noxious stimuli. *C. elegans* genome sequencing revealed the presence of several genes encoding TRP ion channel proteins with important sequence homologies to mammalian TRP channels including TRPVs [3]. Specifically, seven TRP subfamilies including TRPV orthologs (e.g. OSM-9 and OCR-1-4) were characterized. Furthermore, it has been established that *C. elegans* TRP channels are associated with behavioral and physiological processes, including sensory transduction [4, 5]. Many *C. elegans* TRP channels share similar activation and regulatory mechanisms with higher species including mammals. Interestingly, our recent paper revealed that capsaicin can impede nocifensive response of *C. elegans* to noxious heat (i.e. 32°C – 35°C) following a sustained exposure and the effect was reversed 6h post capsaicin exposure [6]. Furthermore, capsaicin’s target was the *C. elegans* transient receptor potential channel OCR-2 and not OSM-9. Additional experiments showed anti-nociceptive effect for other capsaicin analogs, including olvanil, gingerol, shogaol and curcumin.

*In vivo* animal testing is required early during the discovery phase in order to estimate some critical pharmacological parameters (e.g., volume of distribution, clearance, half-life, and bioavailability) and evaluate efficacy and toxicity. However, in pain research, it is almost impossible to screen a complete compound library using a validated animal model of pain due to numerous limitations (e.g. ethical considerations, time limitation to allow the development of the pathological conditions, technical challenges associate with behavior evaluations and all associated cost). These limitations are an import burden for the development of new analgesics. The development of new analgesic drugs is becoming a priority due to the side effects of marketed drugs and a notable increase in the number of patients suffering from chronic pain [7, 8]. It is therefore essential to improve our capacity to test potential therapeutic effects on a larger number of molecules during the early discovery phase in order to better rank compounds for pre-clinical development, and therefore, significantly improve the success rate and eventually bring better drugs to relief patients. One long-term goal is to perform compound library synthesis using the vanillyl group as a primary template and leverage our core knowledge on TRPV1 ligands. Ligand-receptor interactions are classically associated with the pharmacophore features and the vanillyl group has a fundamental role in vanilloids specific interaction with the TRPV1 [9, 10]. Though, it is important to validate if *C. elegans* can be used as a predictive tool of drug efficacy and ultimately improve the animal-to-human translational success.

Eugenol (4-allyl-2-methoxyphenol) is a vanilloids and it is one of the major constituents of clove oil (*Eugenia aromatic*) [11]. Eugenol was frequently used in dentistry as a local analgesic [12], moreover antibacterial [13] and neuroprotective effects [14] were also reported. The biphasic pharmacological effect of eugenol includes an initial pungent effect followed by a delayed analgesia and its chemical structure similar to capsaicin suggest that both compounds are interacting with the same receptor, the transient receptor potential vanilloid 1 (TRPV1) in mammals [15]. Capsaicin and other vanilloids including eugenol pungent action is associated with the initial activation of the TRPV1 evoking membrane depolarization followed by the generation of action potential leading to heat and pain sensation. However, following sustained stimulation, TRPV1 agonists will elicit receptor desensitization, leading to alleviation of pain, a consequence of receptor conformational changes [37]. In recent years, we revealed that vanilloids can effectively alleviate pain in several animal models [16, 17, 18, 19, 20].

Our hypothesis is that heat avoidance behavior is regulated by TRPV ortholog channels in *C. elegans* and the exposure to eugenol and other vanilloids will hamper nocifensive response of *C. elegans* to noxious heat. The objective of this study is to characterize the targeted vanilloid exposure–response relationships using *C. elegans* and heat avoidance behavior analysis [21]. Selected vanilloids and known TRPV1 ligands that will be tested are displayed in Fig. 1. This study is important in order to demonstrate the adequacy of the experimental model.

**Figure 1.**
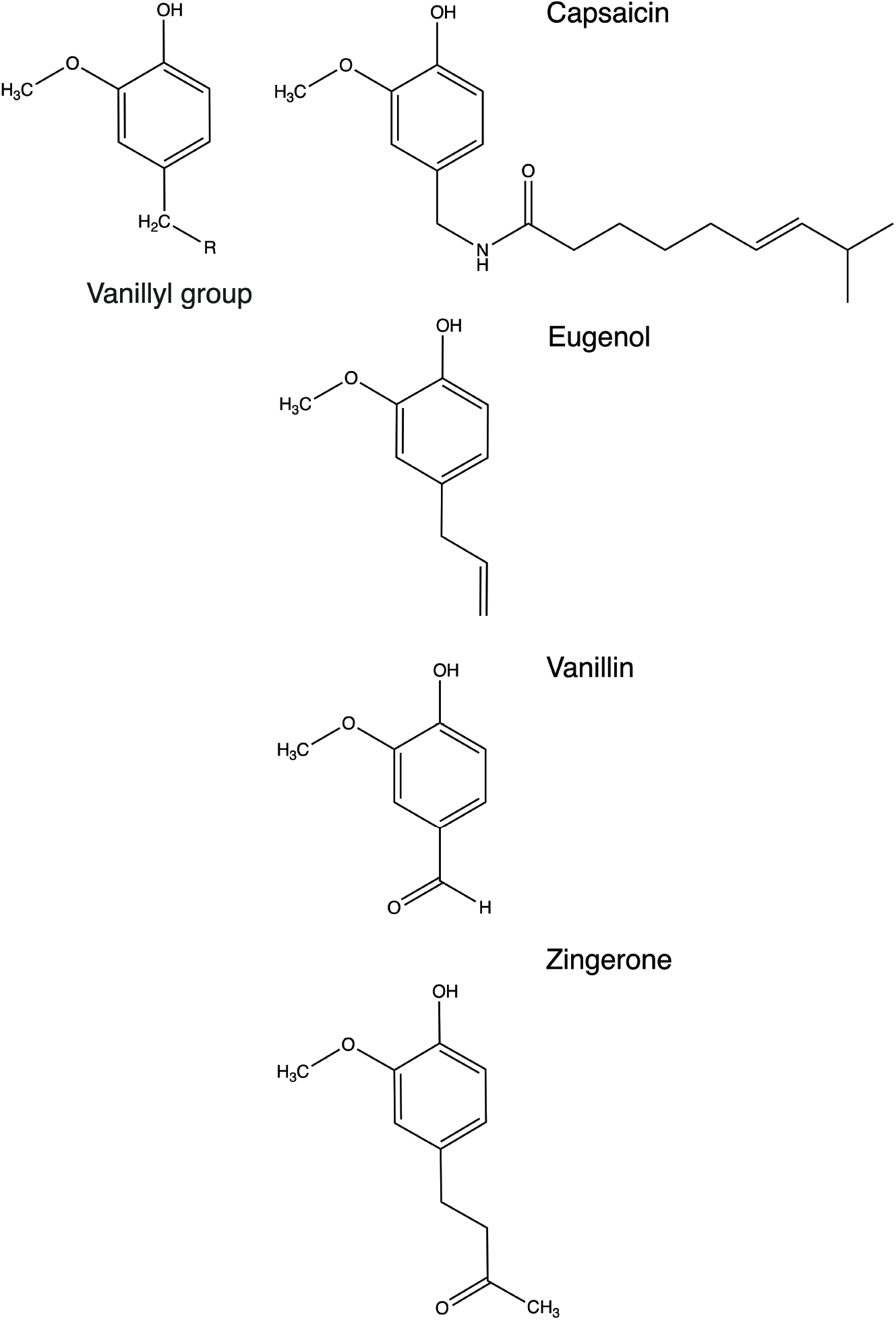
Molecular structure of known TRPV1 ligands capsaicin, eugenol, vanillin and zingerone. Ligand-receptor interactions are characteristically associated with the pharmacophore features and the vanillyl group play a fundamental role in interaction with the TRPV1.

## Materials and Methods

### Chemicals and reagents

All chemicals and reagents were obtained from Fisher Scientific (Fair Lawn, NJ, USA) or MilliporeSigma (St-Louis, MO, USA). Eugenol, vanillin and zingerone were purchased from Toronto Research Chemicals (North York, ON, CAN).

### *C. elegans* strains

The N2 (Bristol) isolate of *C. elegans* was used as a reference strain. Mutant strains used in this study included: *ocr-1* (ak46), *ocr-2* (yz5), *ocr-3* (ok1559)*, ocr-4* (vs137) and *osm-9* (yz6). N2 (Bristol) and other strains were obtained from the Caenorhabditis Genetics Center (CGC), University of Minnesota (Minneapolis, MN, USA). Strains were maintained and manipulated under standard conditions as described [22, 23]. Nematodes were grown and kept on Nematode Growth Medium (NGM) agar at 22°C in a Thermo Scientific Heratherm refrigerated incubator. Experiments and analyses were performed at room temperature unless otherwise noted.

### *C. elegans* pharmacological manipulations

Each vanilloid (i.e. eugenol, vanillin and zingerone) was dissolved in Type 1 Ultrapure Water at a concentration of 25 μM. The solution was warmed for brief periods combined with vortexing and sonication for several minutes to completely dissolve the compound. Then, dilution at 10 μM, 2 μM, 1 μM and 0.5 μM in Type 1 Ultrapure Water was performed by serial dilution for each compound. *C. elegans* were isolated and washed according to protocol outline by Margie *et al*. [23]. After 72 hours of feeding and growing on 92 × 16 mm petri dishes with NGM, the nematodes were exposed to vanilloids solutions. An aliquot of 7 mL of vanilloid solutions was added producing a 2-3 mm solution film (solution is partly absorbed by NGM), therefore, nematodes were swimming in solution. *C. elegans* were exposed for specific times, isolated and washed thoroughly prior behavior experiments. To test for the remanent effect (i.e. 6h latency), after exposure to solutions of eugenol, vanillin or zingerone, nematodes were isolated, carefully washed and deposit on NGM free of vanilloids. The NGM agar free of vanilloids with nematodes was kept at 22°C in an incubator for 6h (i.e. remanent effect/latency test) and thermal avoidance response was retested.

### Thermal avoidance assay

The method we proposed in this manuscript for the evaluation of thermal avoidance was modified from the four quadrants strategies previously described [23, 24] and used in previous successfully published work [6, 21, 25]. The experimental schematics are illustrated in supplementary Figure S1. Briefly, experiments were performed on 92 × 16 mm petri dishes divided into four quadrants. A middle circle delimited (i.e. 1 cm diameter) an area where *C. elegans* were not considered. Petri dishes were divided into quadrants; two stimulus areas (A and D) and two control areas (B and C). Sodium azide (i.e. 0.5M) was used in all quadrants to paralyze the nematodes. Noxious heat was created with an electronically heated metal tip (0.8 mm diameter) producing a radial temperature gradient (e.g. 32-35°C on the NGM agar at 2 mm from the tip measured with an infrared thermometer). Nematodes were isolated and washed according to a protocol outlined by Margie *et al.* [23]. The nematodes tested were off food during all experimentations. The nematodes (i.e. 100 to 300 young adult worms) were placed at the center of a marked petri dish and after 30 minutes, they were counted per quadrant. Note that nematodes that did not cross the inner circle were not considered. The Thermal avoidance Index (TI) formula is shown in supplementary Figure S1. Both TI and the animal avoidance percentage were used to phenotype each tested *C. elegans* experimental groups. The stimulus temperature used was based on previous experiments [2, 6, 21, 25].

### Statistical analysis

Behavior data were analyzed using a one-way ANOVA followed by Dunnett multiple comparison test (e.g. WT(N2) was the control group used). For eugenol and its analogs, the data presented in Fig 4B were analyzed using a two-tailed Student’s t-test (pairwise comparison). Significance was set a priori to p < 0.05. The statistical analyses were performed using PRISM (version 8.3).

### Molecular modeling

The property calculations were performed using Spartan 18 software Wavefunction, Inc. Irvine, CA, USA [26]. Primary, the 3D space-filling models (i.e. CPK models) of capsaicin, eugenol, vanillin and zingerone were generated. Then, a systematic conformational search and analysis were performed to determine the more stable conformers for each compound, presenting the minima energy. Thus, the lowest energy conformer was obtained using MMFF molecular mechanics models by refining the geometry for each studied molecule. Using these structures optimized, calculations of molecular properties and topological descriptors were performed using density functional method [27], software algorithms hybrid B3LYP model [27, 28, 29] and polarization basis set 6-31G* [26, 30] in a vacuum for equilibrium geometry at ground state.

## Results and discussion

In our recent study, we have shown for the first-time, the anti-nociceptive effect of capsaicin in *C. elegans* following a controlled and prolonged exposure [6]. We also identified the capsaicin target, OCR-2. Additionally, other capsaicin analogs displayed similar anti-nociceptive effects. The use of capsaicin in patients is limited due to its pungent action creating a temporal intense pain sensation [31]. Interestingly, eugenol’s pungent action is significantly less compared to capsaicin [15]. This difference was associated with the smaller aliphatic chain at the C4 position of the vanillyl group. Thus, we wanted to verify if vanilloids with a shorter aliphatic chain at the C4 position of the vanillyl group conserve the anti-nociceptive effect in *C. elegans* while the pungent effect is reduced. This is particularly important in the spirit of drug discovery.

The thermal avoidance assay performed is described in supplementary Figure S1 and was specifically used to assess if eugenol, vanillin or zingerone (e.g. selected vanilloids with a shorter aliphatic chain at the C4 position) could hamper nocifensive response to noxious heat. The first experiment included an assessment of the mobility and bias of WT (N2) and mutants *ocr-1*, *ocr-2*, *ocr-3, ocr-4* and *osm-9* nematodes in absence or in presence of eugenol, vanillin or zingerone. As shown in Figure 2, no quadrant selection bias was observed for all *C. elegans* experimental groups or genotypes tested with or without eugenol, vanillin or zingerone exposure (25 μM). Results showed that nematodes were not selecting any specific quadrant and were uniformly distributed after 30 minutes following the initial nematode deposition at the center of the petri dish.

**Figure 2.**
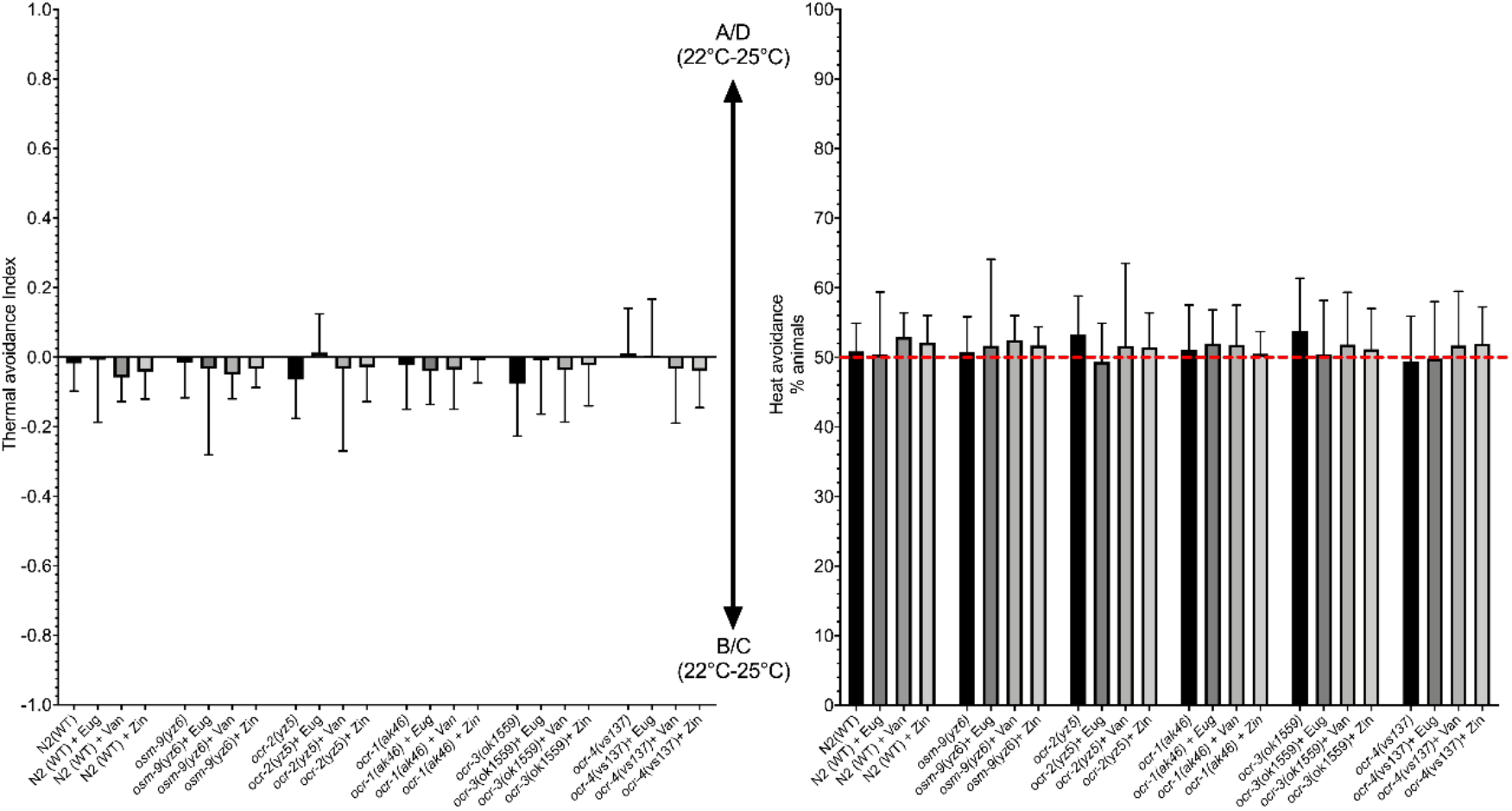
Comparison of the mobility and bias for WT (N2) and mutants *ocr-1*, *ocr-2*, *ocr-3*, *ocr-4* and *osm-9* nematodes in plates divided into quadrants conserved at constant temperature (22°C) and no stimulus was applied (negative control). No quadrant selection bias was observed for all *C. elegans* genotype tested in absence or presence of eugenol (Eug), vanillin (Van) or zingerone (Zin) at 25 μM.

After a prolonged stimulation, TRPV1 agonists elicit receptor desensitization, leading to alleviation of pain (or anti-nociceptive effect in *C. elegans*), which results from conformational changes, along with subsequent decrease of the release of pro-inflammatory molecules and neurotransmitters following exposures to noxious stimuli [9, 10, 32]. Therefore, we have exposed nematodes to eugenol (Figure 3A), vanillin (Figure 3B) or zingerone (Figure 3C) in solution and consequently had complete control of time and exposure levels. As displayed in Figure 3, data revealed a dose–response relationship with a significant anti-nociceptive effect following a 1h exposure to all 3 vanilloids at concentrations ranging from 0.5 μM to 25 μM when compared with the WT (CTL) group. Significant differences in response were observed only for eugenol and zingerone as shown in Figure 3A and Figure 3C. Following eugenol, vanillin or zingerone exposition, nematodes were carefully washed and transferred on NGM agar kept at 22°C in an incubator for 6h (i.e. residual effect/latency test). Then, the thermal avoidance response was retested. As shown in Figure 3, results suggest that after 6h post exposure to all 3 vanilloids, *C. elegans* thermal avoidance response returned to normal. Thus, no residual anti-nociceptive effects of eugenol, vanillin or zingerone were observed after 6h. Eugenol, vanillin or zingerone initial sustained exposure is a key factor to observe vanilloid receptor desensitization as we have previously demonstrated for capsaicin [6].

**Figure 3.**
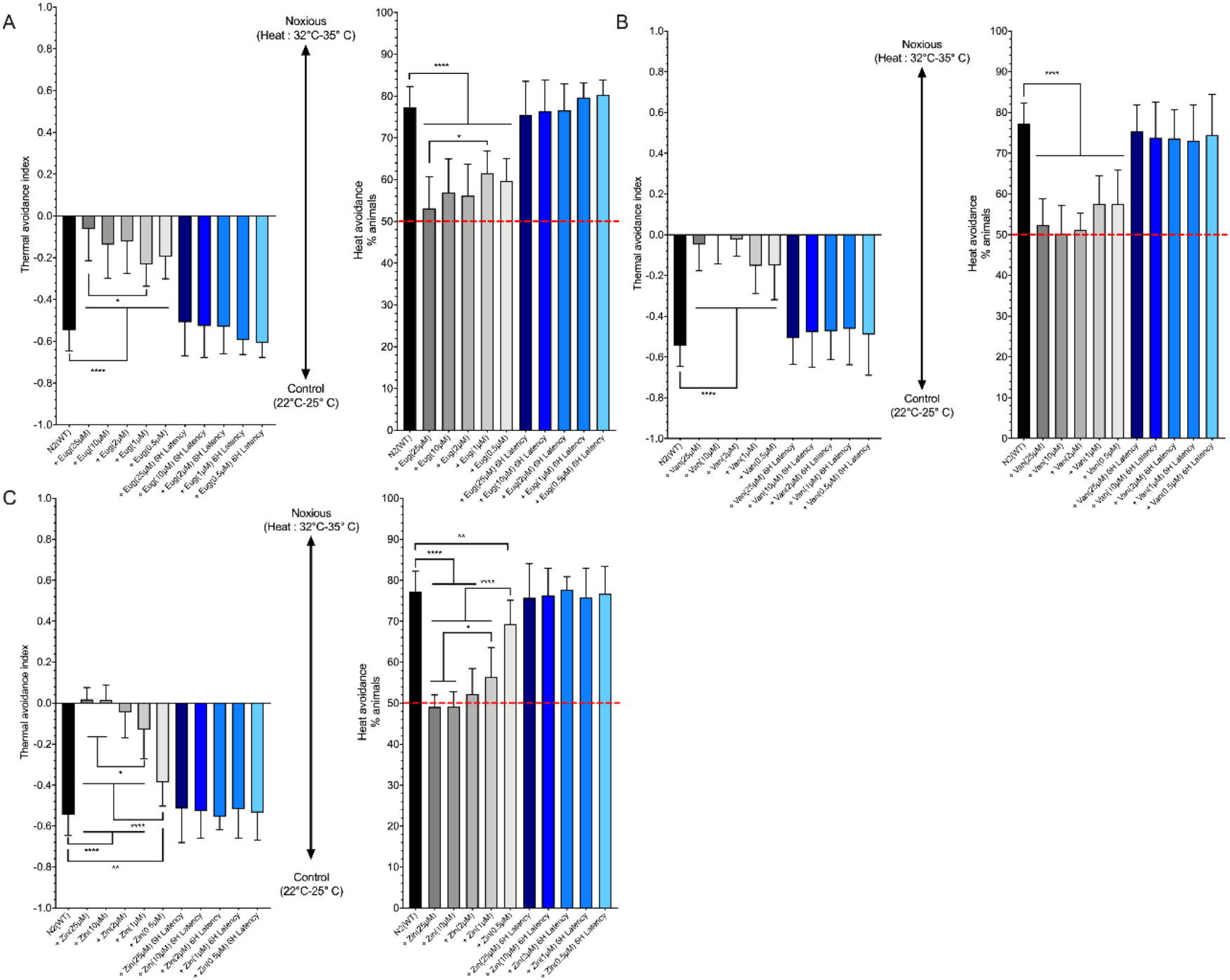
Assessment of the pharmacological effect of vanilloids on thermal avoidance in *C. elegans*. Nematodes were exposed to eugenol, vanillin or zingerone for 60 min prior behavior experimentations. **A)** Eugenol dose-response assessment. **B)** Vanillin dose-response assessment. **C)** Zingerone dose-response assessment. Antinociceptive effect observed following *C. elegans* exposure to eugenol, vanillin or zingerone was dose dependent. Additionally, 6h post exposition, nociceptive behavior returns to normal for all doses and molecules tested. **** p < 0.0001, ** p < 0.01, *p < 0.05 (ANOVA - Tukey’s multiple comparisons test).

Exposure–response relationship can be rationalized when we characterize the relationship involved in the target engagement of the tested molecule. Pharmacological effect can be measured when the tested molecule binds to a sufficient fraction of the target leading to a measurable response (i.e. physiological or phenotypic changes). Thus, experiments were conducted on specific *C. elegans* mutants (i.e. *ocr-1*, *ocr-2*, *ocr-3*, *ocr-4* and *osm-9*) to potentially identify eugenol, vanillin or zingerone target receptors. *C. elegans* mutants were exposed to eugenol, vanillin or zingerone concentration of 25 μM for 60 min prior behavior experiments. As seen in Figure 4A, all mutants, except *ocr-4* (vs137) appeared less sensitive to noxious heat compared with the WT (N2) group. However, they are still sensitive and may suggest redundancy in receptor function. There is no clear evidence eugenol (Figure 4B) or vanillin (Figure 4C) target a specific vanilloid ortholog receptors (e.g. OCR2 or OSM-9) since all tested mutants nocifensive responses to noxious heat were impeded following exposure to both molecules. The amplitude might be less important in *ocr-2* mutant but no clear conclusion can be made based on these experiments. However, no significant zingerone effect (p > 0.05) was observed in *ocr-2* mutant (Figure 4D) suggesting that zingerone targets ORC-2, a transient receptor potential channel, vanilloid subfamily and a mammalian capsaicin receptor-like channel as reported for capsaicin [6]. Thermal nociceptive neurons express heat- and capsaicin-sensitive TRPV channels, OCR-2 and OSM-9 are most likely the receptors targeted by vanilloids. These data sets clearly demonstrate anti-nociceptive effects of eugenol, vanillin and zingerone in *C. elegans*. The data shown in Figure 5 compared anti-nociceptive effect of capsaicin with eugenol, vanillin or zingerone at high concentration (i.e 10μM) and low concentration (i.e. 2μM). As shown, the difference is wider at low concentration (Figure 5B) suggesting significantly more pronounced anti-nociceptive effect for vanillin and zingerone compared with capsaicin. Eugenol’s effect appeared more similar with capsaicin but still showed a more pronounced anti-nociceptive effect at low concentration (i.e. 2μM). These results are interesting since eugenol, vanillin and zingerone are significantly less pungent compared to capsaicin.

**Figure 4.**
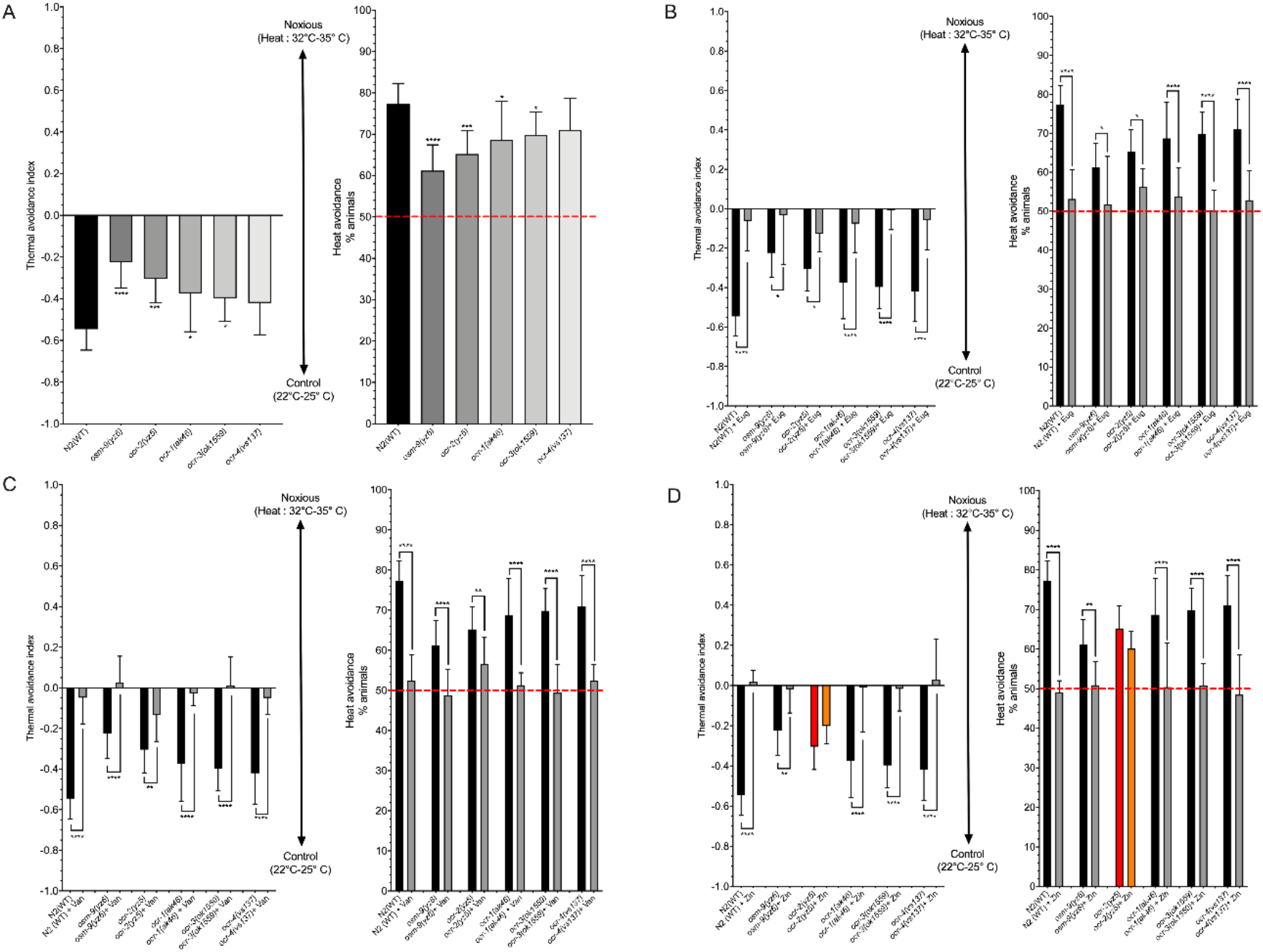
Identification of TRPV orthologs responsible for the anti-nociceptive effect observed for eugenol, vanillin and zingerone. **A)** Heat avoidance behavior phenotyping of tested mutants. Heat avoidance is partly impaired in each mutant, except for *ocr-4*(vs137). **** p < 0.0001, *** p < 0.001, *p < 0.05 (ANOVA - Dunnett’s multiple comparison test) **B)** Eugenol-induced anti-nociceptive effect. **C)** Vanillin-induced anti-nociceptive effect. **D)** Zingerone-induced anti-nociceptive effect. Similarly, to capsaicin (Nkambeu et al. 2020) the data suggest that zingerone exert is anti-nociceptive effects through the ORC-2 *C. elegans* TRPV ortholog. However, results for eugenol and vanillin may indicate redundancy in receptor involvement. **** p < 0.0001, ** p < 0.01, *p < 0.05 (ANOVA - Sidak’s multiple comparisons test).

**Figure 5.**
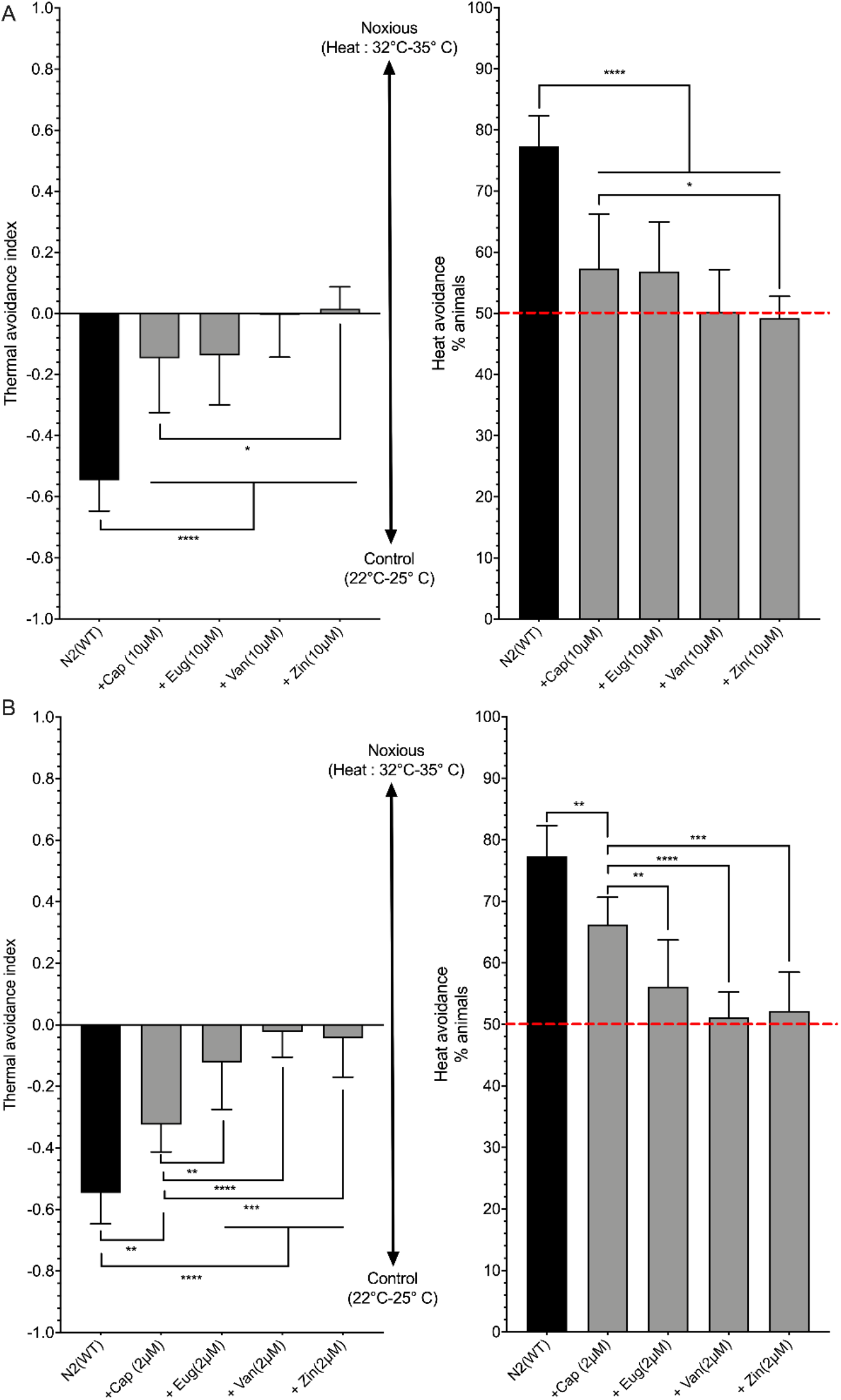
Assessment of the anti-nociceptive and desensitizing effects of vanilloid analogs of capsaicin (Cap). All molecules compared at 10 μM and 2μM produced significant anti-nociceptive and desensitizing effects in *C. elegans*. Moreover, eugenol, vanillin and zingerone generate significantly more anti-nociceptive and desensitizing effects in *C. elegans* compared to capsaicin. **** p < 0.0001, *** p < 0.001 ** p < 0.01, *p < 0.05 (ANOVA - Tukey’s multiple comparisons test).

Further structural and physicochemical analyses were performed. Key parameters for quantitative structure-property relationships (QSPR) and quantitative structure-activity relationships (QSAR) analysis are reported in Table 1. It was previously established [33], in order to increase success of drug candidates, that compounds should respect the following conditions: 1) maximum five hydrogen bond donors (as the total number of nitrogen-hydrogen and oxygen-hydrogen bonds); 2) maximum 10 hydrogen bond acceptors (as the total number of nitrogen and oxygen atoms); 3) the molecular weight should be lower than 500 Da; 4) the octanol-water partition coefficient (log P) value must be less than 5. Table 1 showed the calculated molecular properties from CPK and from Wavefunction models for capsaicin, eugenol, vanillin and zingerone obtained for the most stable conformer of each after geometry minimization: dipole moment, ovality, polarizability, the log P, the number of hydrogen bond donors (HBDs) and acceptor sites (HBAs), area, volume, polar surface area (PSA) and energies of frontier molecular orbitals (FMOs). Area and volume values are in the same ascending order then the respective molecular weights and increase in the following order: vanillin < eugenol < zingerone < capsaicin. However, PSA is not correlating with molecular weight order specifically for vanillin. PSA is associated with the interaction topography and potential steric hindrance [34]. A recent study suggests the PSA is a key descriptor to demystify relative biological activity in a drug library composed of structurally related analogs [35]. Interestingly, the ratio between PSA and molecular weight paralleled with the observed relative anti-nociceptive effect are shown in Figure 5. However, the correlation between the PSA values or the ratio of the PSA and molecular weight values and the anti-nociceptive effect based on the current-set of molecules is not possible and would require a much larger training-set to be statistically relevant. The ovality index characterizes the deviation from the spherical form, considering its value of 1 for an ideal spherical shape. From our calculations we observed the following variation of this parameter: 1.61 (capsaicin) < 1.37 (zingerone) < 1.32 (eugenol) < 1.26 (vanillin). The ovality index is associated with molecular surface area and Van der Waals volume, and it increases with the increase of structural linearity explaining why capsaicin has the highest value. It is also an important parameter for potential steric hindrance and receptor binding specificity. Interestingly, as revealed in our previous study [6] and in Figure 4, only capsaicin and zingerone appear to specifically interact with OCR-2 while eugenol and vanillin are not interacting preferably with OCR-2. Capsaicin and zingerone have higher ovality index values compared to eugenol and vanillin. Generally, the log P value is associated with the lipophilicity of a compound and it can be used to predict the drug absorption and distribution. Interestingly, the calculated log P values for capsaicin and eugenol are significantly higher compared to vanillin and zingerone. As shown in Figure 5, despite showing significant anti-nociceptive effect, capsaicin and eugenol have significantly less effect at 2μM compared to vanillin and zingerone. This may indicate that higher lipophilicity can affect the molecule’s ability to reach the target since a more important fraction of the molecules can be sequestered in animal fat compartments or by cell membrane components. All 4 tested molecules had log P < 5, a criterion for a good drug candidate. The molecular frontier orbitals are important descriptors related to the reactivity of molecules and therefore center to explore ligand-receptor interactions [35, 36]. The HOMO energy is linked to the tendency of a molecule to donate electrons to empty molecular orbitals of partner molecules. Conversely, the LUMO energy indicates the ability to accept electrons. The frontier molecular orbital density distribution of the studied compounds is shown in Figure 6 (HOMO) and Figure 7 (LUMO) for capsaicin, eugenol, vanillin and zingerone. Red and blue areas relate to positive and negative values of the orbital. Interestingly, some distinct features appear for vanillin and zingerone with the group linked to C4 (para position from the hydroxy group) being able to accept electrons (LUMO) from partner molecules. The energy gap (ΔE = | HOMO-LUMO |) obtained provided insight on relative stability and reactivity of the molecules. Accordingly, the higher value means the molecule is more stable and less reactive. The obtained energy gap increases in the order: eugenol (5.70) > capsaicin (5.67) > zingerone (4.92) and vanillin (4.67). (4.36 < 4.39 < 5.09). These results suggest eugenol presents the lowest reactivity (the most chemically stable) followed by capsaicin, zingerone and vanillin (the most reactive). These results also appear coherent with the relative anti-nociceptive effect shown in Figure 5 keeping in mind the minute difference between capsaicin and eugenol at high concentration (10 μM). However, at low concentration (2μM) eugenol has a more pronounced anti-nociceptive effect compared to capsaicin but eugenol’s lower PSA and ovality, affecting steric hindrance, values may play an important role when comparing both molecules at lower concentration.

**Table 1.** Predicted molecular properties for capsaicin, eugenol, vanillin and zingerone, using DFT method, B3LYP model using the 6-31G* basis set, in vacuum, for equilibrium geometry at ground state.

|  | Capsaicin | Eugenol | Vanillin | Zingerone |
| --- | --- | --- | --- | --- |
| **Molecular properties** |  |  |  |  |
| Formula | C<sub>18</sub>H<sub>27</sub>NO<sub>3</sub> | C<sub>10</sub>H<sub>12</sub>O<sub>2</sub> | C<sub>8</sub>H<sub>8</sub>O<sub>3</sub> | C<sub>11</sub>H<sub>14</sub>O<sub>3</sub> |
| Molecular weight (amu) | 305.418 | 164.204 | 152.149 | 194.230 |
| Energy (au) | -982.596295 | -538.697381 | -535.318126 | -653.261314 |
| Dipole moment (debye) | 1.41 | 2.53 | 2.75 | 2.24 |
| E HOMO (eV) | -5.57 | -5.45 | -6.07 | -5.37 |
| E LUMO (eV) | 0.10 | 0.25 | -1.40 | -0.45 |
| **QSAR properties from CPK model** |  |  |  |  |
| Area (Å<sup>2</sup>) | 382.96 | 207.17 | 175.13 | 232.78 |
| Volume (Å<sup>3</sup>) | 346.53 | 184.38 | 154.14 | 209.37 |
| PSA (Å<sup>2</sup>) | 48.746 | 24.408 | 38.963 | 37.973 |
| Ovality | 1.61 | 1.32 | 1.26 | 1.37 |
| **QSAR properties from computed electron density** |  |  |  |  |
| Log P | 3.66 | 2.57 | 1.27 | 1.95 |
| HBD count | 2 | 1 | 1 | 1 |
| HBA count | 4 | 2 | 3 | 3 |
| Polarizability (10<sup>-30</sup> m<sup>3</sup>) | 68.15 | 54.98 | 52.77 | 57.20 |

**Table 2.**
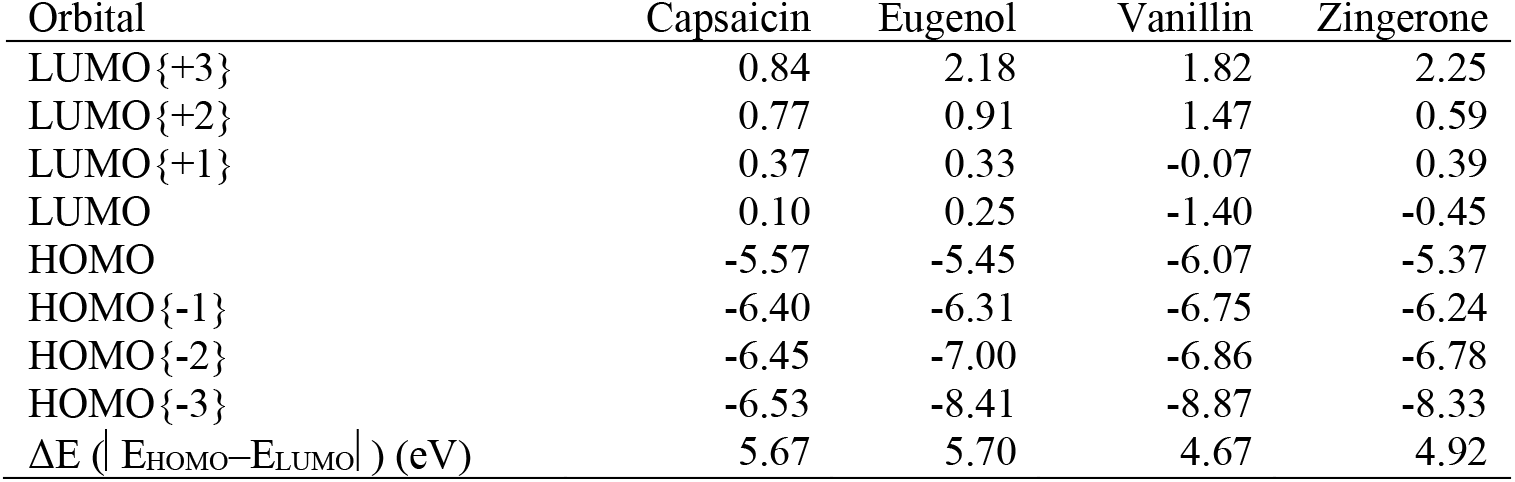
Capsaicin, eugenol, vanillin and zingerone energetic levels (eV) of intermediary molecular orbitals (MO).

**Figure 6.**
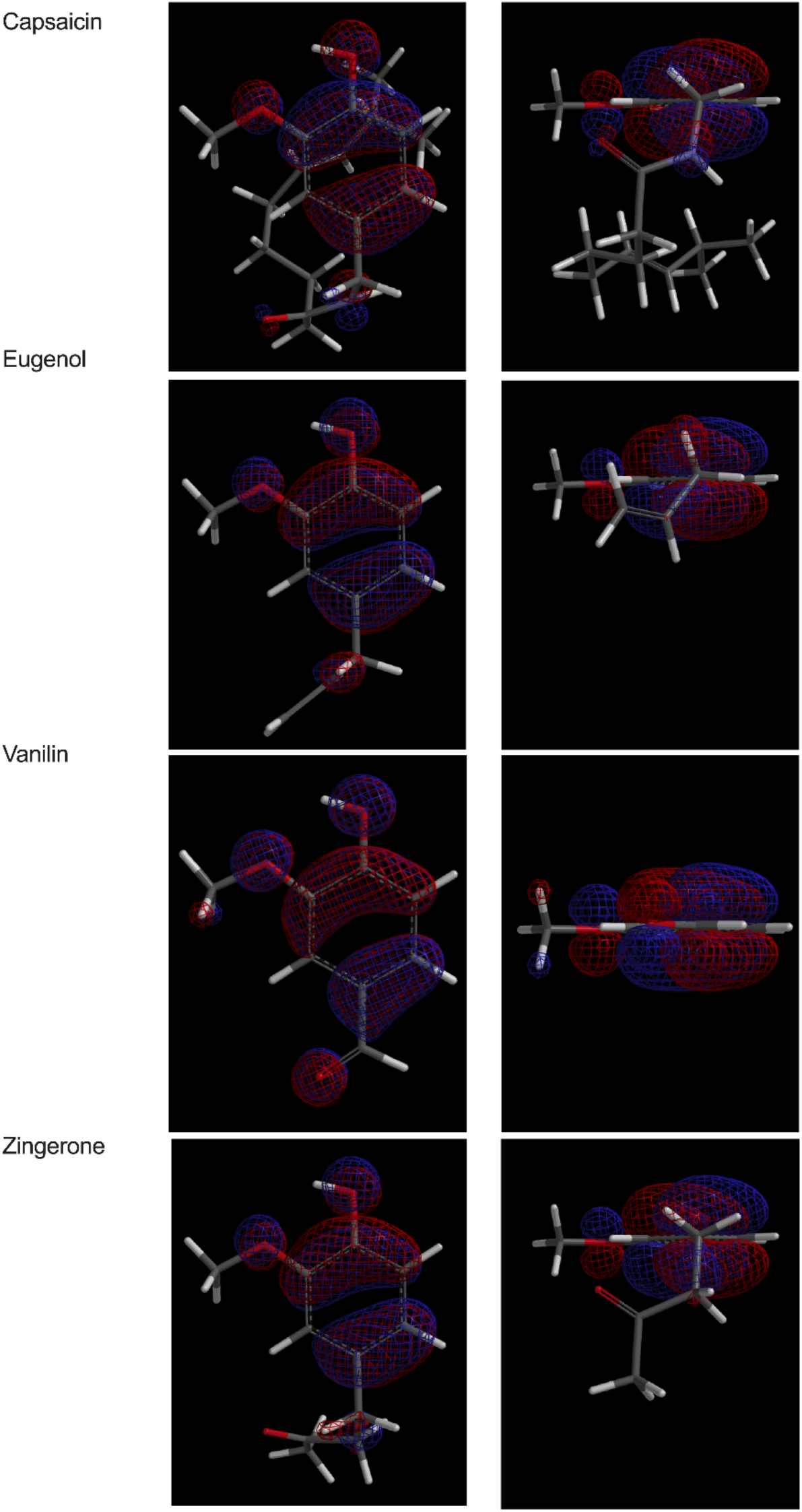
HOMO orbital plots of capsaicin, eugenol, vanillin and zingerone using DFT method, B3LYP model and 6-31G* basis set, in vacuum for equilibrium geometry at ground state. Molecular orbitals provide important indications about chemical reactivity. The orbital drawing identifies regions where the orbital takes on a significant value, either positive (blue) or negative (red).

**Figure 7.**
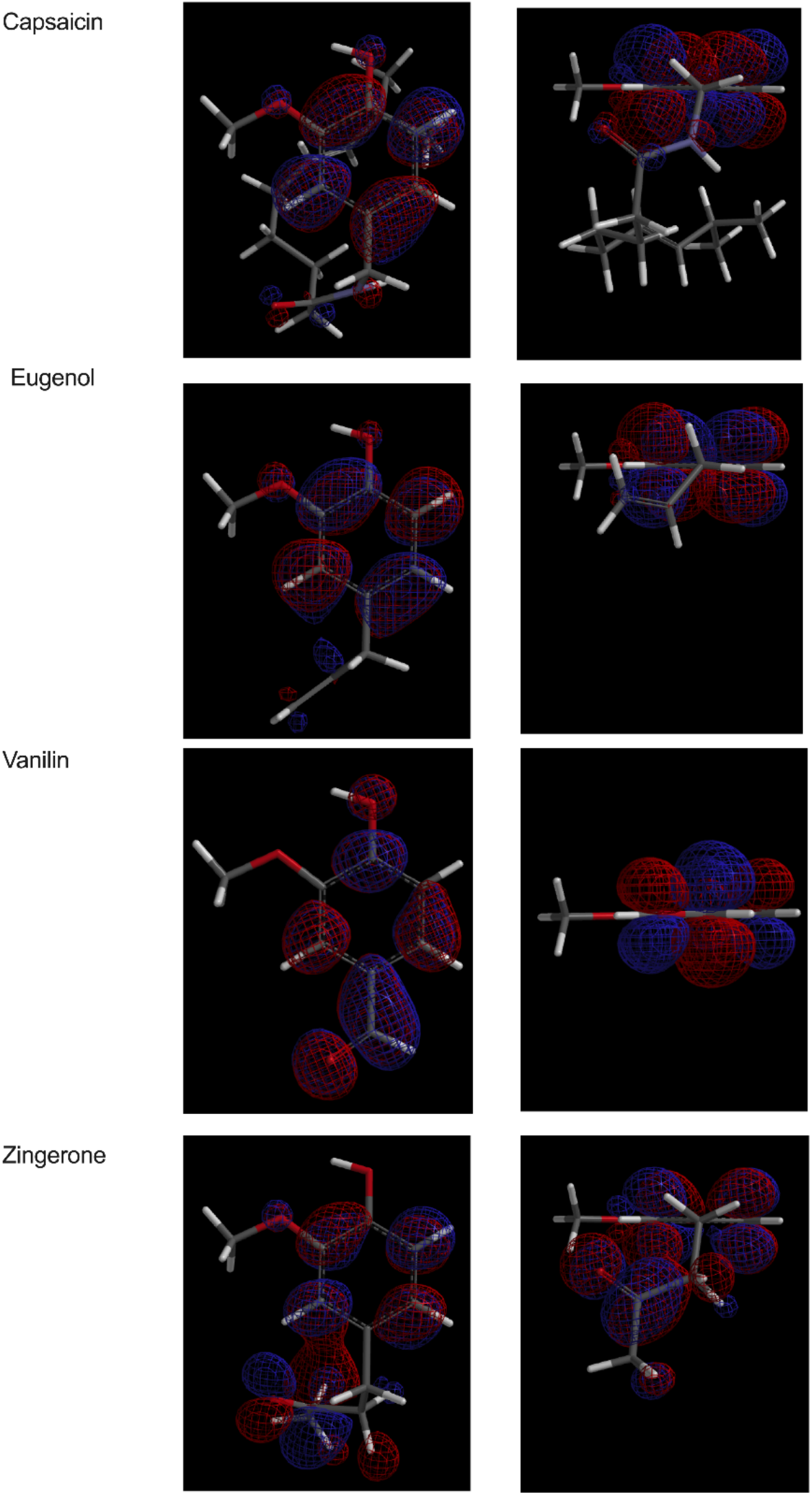
LUMO orbital plots of capsaicin, eugenol, vanillin and zingerone using DFT method, B3LYP model and 6-31G* basis set, in vacuum for equilibrium geometry at ground state. Molecular orbitals provide important indications about chemical reactivity. The orbital drawing identifies regions where the orbital takes on a significant value, either positive (blue) or negative (red).

## Conclusion

This study clearly revealed eugenol’s, vanillin’s and zingerone’s anti-nociceptive effect in *C. elegans* following a controlled and prolonged exposition. These molecules effectively impede nocifensive response to noxious heat. Additionally, eugenol and vanillin did not target specifically the OCR-2 or OSM-9 but zingerone did specifically target the OCR-2 similarly to capsaicin. However, functional redundancy between OCR-2 and OSM-9 is anticipated. As shown, the modeling of QSPR/QSAR properties and molecular frontier orbitals provide new understandings of vanilloids anti-nociceptive effects observed in *C. elegans*. These analyses are key chemoinformatics parameters to build a large vanilloid compound database to design drugs with desired biological properties. *C. elegans* could be used to rank these compounds for further evaluation in experimental models of pain and ultimately improve animal-to-human translational success.

## Acknowledgements

This project was funded by the National Sciences and Engineering Research Council of Canada (F. Beaudry discovery grant no. RGPIN-2015-05071 and RGPIN-2020-05228). Laboratory equipment was funded by the Canadian Foundation for Innovation (CFI) and the *Fonds de Recherche du Québec (FRQ)*, the Government of Quebec (F. Beaudry CFI John R. Evans Leaders grant no. 36706). A PhD scholarship was awarded to J. Ben Salem with a grant obtained from *Fondation de France*.

## References

1. Van Pelt KM, and Truttmann MC (2020) *Caenorhabditis elegans* as model system for studying aging-associated neurodegenerative diseases. Translational Medecine of Aging 4:46 – 72

2. Wittenburg N, Baumeister R (1999) Thermal avoidance in *Caenorhabditis elegans*: An approach to the study of nociception. Proc Natl Acad Sci 96(18):10477–10482

3. Kahn-Kirby AH, Bargmann CI (2006) TRP channel in *C. elegans*. Annual Review of Physiology 68:719 – 736

4. Glauser DA, Chen WC, Agin R, MacInnis B, Hellman AB, Garrity PA, Man-WahTan, Goodman MB (2011) Heat avoidance is regulated by Transient Receptor Potential (TRP) channels and a neuropeptide signaling pathway in *Caenorhabditis elegans*. Genetics Society of America 188:91–103

5. Venkatachalam K, Luo J, Montell C (2014) Evolutionarily Conserved, Multitasking TRP Channels: Lessons from Worms and Flies. Handb Exp Pharmacol 223:937 – 962

6. Nkambeu B, Ben Salem J, Beaudry F (2020) Capsaicin and Its Analogues Impede Nocifensive Response of *Caenorhabditis elegans* to Noxious Heat. Neurochemical Research 45:1851 – 1859

7. Dworkin RH, O’Connor AB, Backonja M, Farrar JT, Finnerup NB, Jensen TS, Kalso EA, Loeser JD, Miaskowski C, Nurmikko TJ, Portenoy RK, Rice ASC, Stacey BC, Treede R-D Turk DC, Wallace MS (2007) Pharmacologic management of neuropathic pain: evidence-based recommendations. Pain 132:237 – 251

8. Borsook D, Hargreaves R, Bountra C, Porreca F (2014) Lost but making progress – Where will new analgesic drugs come from? Science Translational Medecine 6(249):1 – 11

9. Bhattarai S, Tran VH, Duke CC (2001) The stability of Gingerol and Shogaol in aqueous solutions. J Pharm Sci 90(10):1658–1664

10. Zhu X, Li Q, Chang R, Yang D, Song Z, Guo Q, Huang C (2014) Curcumin alleviates neuropathic pain by inhibiting p300/CBP histone acetyltransferase activity-regulated expression of BDNF and Cox-2 in rat model. PLoS ONE 9(3):1 – 9

11. Pramod K, Ansari SH, Ali J (2010) Eugenol: A Natural Compound with Versatile Pharmacological Actions. Natural Product Communications 5(12):1999 – 2006

12. Ohkubo T, Shibata M (1997) The selective capsaicin antagonist capsazepine abolishes the antinociceptive action of eugenol and guaiacol. J Dent Res 76:848 – 851

13. Laekeman GM, Van Hoof L, Haemers A, Van Berghe DA, Herman AG, Vlietink AK (1990) Eugenol, a valuable compound for *in-vitro* experimental research and worthwile for further *in-vivo* investigation. Phytother. Res 4:90 – 96

14. Wie MB, Won MH, Lee KH, Shin JH, Lee JC, Suh HW, Song DK, Kim YH (1997) Eugenol protects neuronal cells from excitotoxic and oxidative injury in primary cortical cultures. Neurosci Lett 225:93 – 96

15. Yang BH, Piao ZG, Kim YB, Lee CH, Lee JK, Park K, Kim JS, Oh SB (2003) Activation of vanilloid receptor 1 (VR1) by eugenol. J Dent Res 82:785 – 785

16. Gauthier ML, Beaudry F, Vachon P (2013) Phytotherapy Research 27(8):125 – 1254

17. Ferland CE, Beaudry F, Vachon P (2012) Antinociceptive Effects of Eugenol Evaluated in a Monoiodoacetate-induced Osteoarthris Rat Model. Phytother Res 26:1278 – 1285

18. Beaudry F, Ross A, Lema, PP, Vachon P (2010) Pharmacokinetics of Vanillin and its Effects on Mechanical Hypersensitivity in Rat Model of Neuropathic Pain. Phytother Res 24:525 – 530

19. Lionnet L, Beaudry F, Vachon P (2010) Intrathecal eugenol administration alleviates neuropathic pain in male Sprague-Dawley rats. Phytotherapy Research 24:1645 – 1653

20. Guénette S, Ross A, Marier J, Beaudry D, Vachon P (2007) Pharmacokinetics of eugenol and its effects on thermal hypersensitivity in rats. European journal of pharmacology 562(1-2):60 – 67

21. Nkambeu B, Ben Salem J, Leonelli S, Amin Marashi F, Beaudry F (2019) EGL-3 and EGL-21 are required to trigger nocifensive response of *Caenorhabditis elegans* to noxious heat. Neuropeptides 73:41–48

22. Brenner S (1974) The genetics of *Caenorhabditis elegans*. Genetics 77:71 – 94

23. Margie O, Palmer C, Chin-Sang I (2013) *C. elegans* Chemotaxis Assay. J Vis Exp 74:1 – 6

24. Porta-De-La-Riva M, Fontrodona L, Villanueva A, Cerôn J (2012) Basic *Caenorhabditis elegans* methods: synchronization and observation. J. Vis. Exp 64:1 – 9

25. Leonnelli S, Nkambeu B, Beaudry F (2020) Impaired Eat-4 Vesicular Glutamate Transporter Leads to Defective Nocifensive Response of Caenorhabditis elegans to Noxious Heat. Neurochemical Research 45:882 – 890

26. Shao Y, Molnar F, Jung Y, Kussmann J, Ochsenfeld C, Brown ST, Gilbert ATB, Slipchenko LV, Levchenko SV, O’Neill DP, DiStasio RA, Lochan RC, Wang T, Beran GJO, Besley NA, Herbert JM, Lin CY, Van Voorhis T, Chien SH, Sodt A, Steele RP, Rassolov VA, Maslen PE, Korambath PP, Adamson RD, Austin B, Baker J, Byrd EFC, Dachsel H, Doerksen RJ, Dreuw A, Dunietz BD, Dutoi D, Furlani TR, Gwaltney SR, Heyden A, Hirata S, Hsu CP, Kedziora G, Khalliulin RZ, Klunzinger P, Lee AM, Lee MS, Liang WZ, Lotan I, Nair N, Peters B, Proynov EI, Pieniazek PA, Rhee YM, Ritchie J, Rosta E, Sherrill CD, Simmonett AC, Subotnik, JE, Woodcock HL, Zhang W, Bell AT, Chakraborty AK, Chipman DM, Keil FJ, Warshel A, Hehre WJ, Schaefer HF, Kong J, Krylov AI, Pmw G, Head-Gordon M (2006) Advances in methods and algorithms in a modern quantum chemistry program pack-age. Physical Chemistry Chemical Physics 8(27):3172 – 3191

27. Parr RG, Yang W (1980) Density Functional Theory of Atoms and Molecules. Oxford : Oxford University Press. International Academy of Quantum Molecular Science vol 3. Springer, Dordrecht

28. Lee C,Yang W, Parr RG (1988) Development of the Colle-Salvetti correlation-energy formula into a functional of the electron density. Physical Review B 37:785 – 789

29. Becke AD (1993) Density functional thermochemistry. III. The role of exact exchange. The Journal of Chemical Physics 98:5648 – 5652

30. Hehre WJ (2003) A Guide to Molecular Mechanics and Quantum Chemical Calculations. Irvine, CA 92612: Wavefunction Inc 1 – 54

31. Saljoughian M (2009) Capsaicin : Risks and Benefits. US Pharm 34(7):HS-17 – HS-18

32. Jara-Oseguera A, Simon SA, Rosenbaum T (2008) TRPV1 : On the road to pain relief. Curr Mol Pharmacol, 1(3):255 – 269

33. Lipinski CA, Lombardo F, Dominy BW, Feeney PJ (2001) Experimental and computational approaches to estimate solubility and permeability in drug discovery and development settings. Advanced Drug Delivery Reviews 46:3 – 26.

34. Schaftenaar G, De Vlieg J (2012) Quantum mechanical polar surface area. J Comput Aided Mol Des 26(3):311 – 318. doi:10.1007/s10822-012-9557-y

35. Clark DE (2011) What has polar surface area ever done for drug discovery? Future Med Chem, 3(4):469 – 484

36. Kizilcan DS, Türkmenoglu B, Güzel Y (2020) The use of the Klopman index as a new descriptor for pharmacophore analysis on strong aromatase inhibitor flavonoids against estrogen-dependant breast cancer. Structural Chemistry 31:1339 – 1351s

37. Jancso, G., Dux, M., Oszlacs, O., & Santha, P. (2008) Activation of the transient receptor potential vanilloid-1 (TRPV1) channel opens the gate for pain relief. British Journal of Pharmacology, 155, 1139 – 1141.

